# G307S DNAM-1 mutation exacerbates autoimmune encephalomyelitis via enhancing CD4^+^ T cell activation

**DOI:** 10.1101/2022.06.08.495273

**Authors:** Rikito Murata, Shota Kinoshita, Kenshiro Matsuda, Atsushi Kawaguchi, Akira Shibuya, Kazuko Shibuya

## Abstract

Although rs763361, which causes a non-synonymous glycine-to-serine mutation at residue 307 (G307S mutation) of the DNAM-1 immunoreceptor, is a single-nucleotide polymorphism (SNP) associated with autoimmune disease susceptibility, little is known about how the SNP is involved in pathogenesis. Here, we established CD4^+^ T cells expressing wild-type or G307S DNAM-1 and showed that the costimulatory signal from G307S DNAM-1 induced greater pro-inflammatory cytokine production and cell proliferation than that from wild-type DNAM-1. The G307S mutation also enhanced the recruitment of the tyrosine kinase Lck and augmented tyrosine phosphorylation at residue 322 (Tyr^322^) of DNAM-1. However, conversion of Tyr^322^ of DNAM-1 to phenylalanine (Y322F mutation) canceled the enhanced activation of CD4^+^ T cell transfectants expressing G307S DNAM-1. Adoptive transfer of myelin-antigen-specific CD4^+^ T cells expressing G307S DNAM-1 into mice exacerbated experimental autoimmune encephalomyelitis compared with the transfer of cells expressing wild-type DNAM-1. These findings suggest that rs763361 is a gain-of-function mutation that enhances DNAM-1-mediated costimulatory signaling for proinflammatory responses.

## Introduction

Autoreactive CD4^+^ T cells play a critical role in the pathogenesis of various autoimmune diseases. Activation of CD4^+^ T cells is finely tuned by signals mediated by the TCR, co-stimulatory molecules, and cytokine receptors, resulting in cytokine production, cell proliferation, and T cell differentiation. Single-nucleotide polymorphisms (SNPs) in molecules regulating the effector functions of CD4^+^ T cells are associated with multiple autoimmune disorders (Parkes et al., 2013; Márquez et al., 2018). Co-stimulatory molecules regulate the effector functions of CD4^+^ T cells by enhancing TCR signaling, and they are promising therapeutic targets of neutralizing antibodies (Edner et al., 2020). Although several functional analyses of disease-susceptibility-associated SNPs of co-stimulatory molecules have been reported (Kofler et al., 2011; Lee et al., 2015; Margraf et al., 2015), little is known about their roles in the pathogenesis of autoimmune diseases or the molecular mechanisms by which they augment the pathogenicity of autoreactive CD4^+^ T cells.

DNAM-1 (also known as CD226) is a signal-transducing adhesion molecule that belongs to the immunoglobulin superfamily. DNAM-1 is predominantly expressed on T cells and natural killer (NK) cells in human peripheral blood mononuclear cells (PBMCs) (Shibuya et al., 1996). Upon binding of DNAM-1 to the ligand CD155 (poliovirus receptor or nectin-like protein 5) or CD112 (nectin-2 or poliovirus receptor-related 2) (Bottino et al., 2003; Tahara-Hanaoka et al., 2004), the tyrosine at residue 322 (Tyr^322^) located in the ITT (immunoreceptor tyrosine tail)-like motif of DNAM-1 is phosphorylated by Src-family kinases to induce downstream signaling (Shibuya et al., 1999, 2003; Zhang et al., 2015). DNAM-1 works as a co-stimulatory molecule in CD4^+^ T cells to promote cytokine production, cell proliferation, and T-helper-1 (Th1)-cell differentiation (Shibuya et al., 2003; Gaud et al., 2018). However, DNAM-1 also reduces forkhead box P3 (Foxp3) stability and the function of regulatory T (Treg) cells (Fourcade et al., 2018; Sato et al., 2021). Moreover, several studies have demonstrated that DNAM-1 exacerbates autoimmune diseases in mice (N. Wang et al., 2020; Shapiro et al., 2020; Dardalhon et al., 2005). Several pieces of evidence have demonstrated that one of the SNPs in the *CD226* locus rs763361 is associated with multiple autoimmune diseases, including multiple sclerosis (Hafler et al., 2009; Bai et al., 2020), type 1 diabetes (Todd et al., 2007; Smyth et al., 2008), rheumatoid arthritis (Maiti et al., 2010; Suzuki et al., 2013), systemic lupus erythematosus (Du et al., 2011), neuromyelitis optica (Liu et al., 2012), primary immune thrombocytopenia (S. Wang et al., 2020), juvenile idiopathic arthritis (Reinards et al., 2015), and autoimmune thyroid disease (Hafler et al., 2009; Saevarsdottir et al., 2020). rs763361 is a non-synonymous mutation that substitutes serine (Ser^307^) for glycine (Gly^307^) at residue 307 (G307S) in the cytoplasmic region of DNAM-1. CD4^+^ T cells isolated from rs763361 carriers show greater increases in phosphorylation of downstream signaling mediators and IL-17 production than the cells of non-carriers (Gaud et al., 2018; Šunina et al., 2021). In contrast, Treg cells obtained from rs763361 carriers show less suppression activity than those of non-carriers (Piédavent-Salomon et al., 2015). In addition, rs763361 shows high linkage disequilibrium with several SNPs, including rs727088, which is located in the 3ʹ-untranslated region of the *CD226* locus and is associated with susceptibility to systemic lupus erythematosus and inflammatory bowel disease via the alteration of DNAM-1 expression (Löfgren et al., 2010; Liu et al., 2017; Jostins et al., 2012). However, the way in which rs763361 is associated with multiple autoimmune diseases remains undetermined.

Here, we show that the G307S mutation is a gain-of-function mutation of DNAM-1 that promotes the effector function of conventional T (Tconv) cells through the enhanced phosphorylation of Tyr^322^. Furthermore, we show that the G307S mutation of DNAM-1 exacerbates experimental autoimmune encephalomyelitis (EAE) in mice

## Results

### G307S mutation of DNAM-1 enhances pro-inflammatory cytokine production by Tconv cells

To examine whether G307S DNAM-1 has a functional role in the development of autoimmune diseases, we used flow cytometry to isolate CD4^+^ Tconv cells and CD4^+^ regulatory T (Treg) cells, as defined by the expression of CD25^Lo^ CD127^Hi^ CD4^+^ and CD25^Hi^ CD127^Lo^ CD4^+^, respectively, from the peripheral blood of healthy volunteers (Figure 1A). We transduced these cells with retrovirus expression vectors encoding either wild-type (WT) or G307S DNAM-1 tagged with either FLAG protein at the N-terminus together with green fluorescent protein (GFP) through the internal ribosome entry site (IRES) or with GFP alone (mock) (Figure 1B and C). The CD4^+^ Tconv and Treg cell transfectants expressed comparable levels of FLAG as well as DNAM-1 on their surfaces (Figure 1-figure supplement 1A and B). After stimulation with anti-CD3 and anti-FLAG monoclonal antibodies (mAbs), Tconv cells expressing FLAG-G307S DNAM-1 produced significantly more IFN-γ and TNF than did those expressing FLAG-WT DNAM-1 (Figure 1D and E), whereas IL-17A and IL-4 were undetectable in both transfectants, even after the same stimulation (Figure 1-figure supplement 1C and D). These results indicated that the pro-inflammatory cytokine production enhanced in Tconv cells by FLAG-G307S DNAM-1 was greater than that enhanced by FLAG-WT DNAM-1.

**Figure 1.**
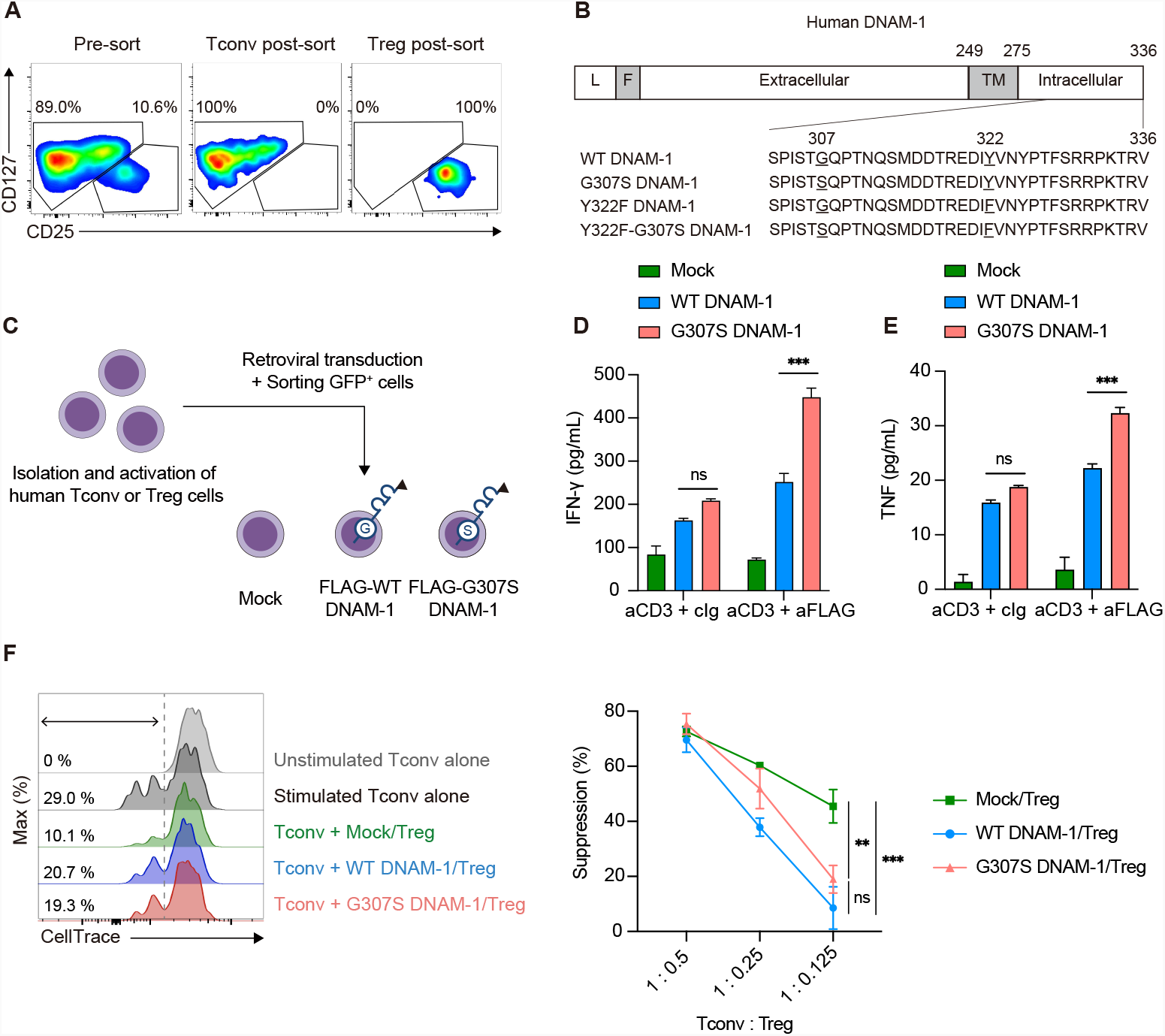
G307S mutation of DNAM-1 enhanced inflammatory cytokine production of Tconv cells. (**A**) Gating strategy for Tconv cells and Treg cells from purified CD4^+^ T cells sorted from human PBMCs. CD4^+^ CD25^Lo^ CD127^Hi^ lymphocytes were sorted as Tconv cells, and CD4^+^ CD25^Hi^ CD127^Lo^ lymphocytes were sorted as Treg cells. (**B**) Schematic diagram of FLAG-DNAM-1. Abbreviation used: L, Leader sequence; F, FLAG; TM, Transmembrane. (**C**) Schematic of generation for FLAG-DNAM-1-expressing human CD4^+^ T cells. Tconv cells and Treg cells from human PBMCs were transduced with a retrovirus encoding Mock (IRES-GFP), FLAG-WT DNAM-1-IRES-GFP, or FLAG-G307S DNAM-1-IRES-GFP, followed by sorting GFP^+^ cells by flow cytometry. (**D and E**) The concentration of IFN-γ and TNF in the culture supernatant of human Tconv cells expressing Mock, FLAG-WT DNAM-1, or FLAG-G307S DNAM-1 after cross-linking with anti-CD3 mAb and either anti-FLAG mAb or control Ig. (n = 3 in each group) (**F**) Human Treg cells expressing Mock, FLAG-WT DNAM-1, or FLAG-G307S DNAM-1 were co-cultured with CellTrace Violet-labeled autologous Tconv cells under the stimulation with anti-CD3 and anti-FLAG mAb. (n = 3 in each group) Data are representative of independent experiments using Tconv cells derived from three different healthy donors (**D and E**) and three independent experiments (**F**). Statistical analysis was performed by using two-way ANOVA with Tukey multiple comparisons test (**D-F**); *P<0.05, **P<0.01, ***P<0.001, ns, not significant.

We next examined whether FLAG-G307S DNAM-1 affected Treg-cell function. Tconv cells were labeled with CellTrace Violet, cocultured with autologous Treg cells expressing FLAG-WT or G307S DNAM-1 or with mock-Treg cells in the presence of beads coated with anti-CD3 and anti-FLAG mAbs, and analyzed by flow cytometry for the proliferation of Tconv cells. There was significantly less proliferation of Tconv cells upon coculture with Treg cells expressing FLAG-WT DNAM-1 than with mock-Treg cells (Figure 1F), suggesting that expression of DNAM-1 decreased Treg-cell function, consistent with our recent finding (Sato et al., 2021). However, the proliferation of Tconv cells upon coculture with Treg cells expressing FLAG-WT was comparable to that upon coculture with cells expressing FLAG-G307S DNAM-1 (Figure 1F). In addition, Foxp3 expression and IL-10 production were equivalent in Treg cells expressing FLAG-WT and those expressing FLAG-G307S DNAM-1 after stimulation with anti-CD3 and anti-FLAG mAbs (Figure 1-figure supplement 1E and F). These findings suggested that G307S DNAM-1 did not affect Treg cell function.

### G307S DNAM-1 enhances tyrosine phosphorylation of DNAM-1

The tyrosine at residue 322 (Tyr^322^) is a potential phosphorylation site involved in DNAM-1-mediated signaling (Shibuya et al., 1999, 2003; Zhang et al., 2015). Considering that the G307S mutation enhanced proinflammatory cytokine production by Tconv cells, we examined the role of this mutation in tyrosine phosphorylation of DNAM-1. We established transfectants of the human T cell line Jurkat stably expressing FLAG-WT DNAM-1, FLAG-G307S DNAM-1, FLAG-Y322F DNAM-1, or FLAG-G307S-Y322F DNAM-1 (Figure 1B, Figure 2-figure supplement 1A). After stimulation of the transfectants with anti-DNAM-1 mAb, FLAG-G307S DNAM-1 showed enhanced tyrosine phosphorylation compared with WT DNAM-1. However, tyrosine phosphorylation was not observed in Y322F DNAM-1 and G307S-Y322F DNAM-1 (Figure 2A). These results suggested that the G307S mutation enhanced the phosphorylation of Tyr^322^ after stimulation with anti-DNAM-1 mAb.

**Figure 2.**
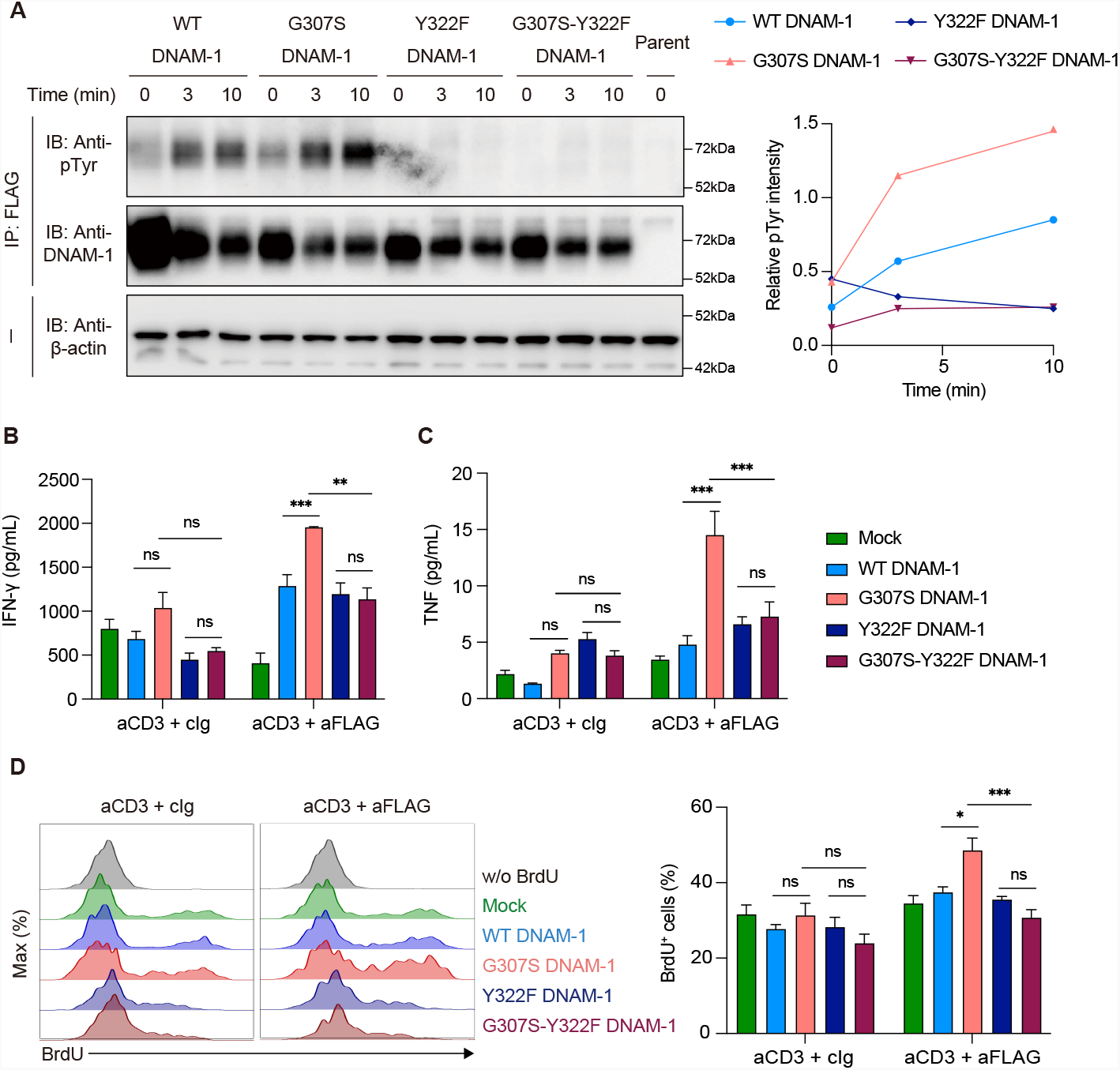
G307S DNAM-1 enhanced tyrosine phosphorylation, inflammatory cytokine production, and proliferation. (**A**) Jurkat T cells expressing FLAG-WT DNAM-1, FLAG-G307S DNAM-1, FLAG-Y322F DNAM-1, or FLAG-G307S-Y322F DNAM-1 were stimulated with anti-DNAM-1 mAb. Cells were immunoprecipitated with anti-FLAG mAb and immunoblotted with anti-phosphotyrosine mAb or anti-DNAM-1. (**B and C**) The concentration of IFN-γ and TNF in the culture supernatant of human Tconv cells expressing Mock, FLAG-WT DNAM-1, FLAG-G307S DNAM-1, FLAG-Y322F DNAM-1, or FLAG-G307S-Y322F DNAM-1 after stimulation with anti-CD3 mAb and either anti-FLAG mAb or control Ig. (n = 3 in each group) (**D**) The BrdU incorporation of human Tconv cells expressing Mock (n = 6), FLAG-WT DNAM-1 (n = 6), FLAG-G307S DNAM-1 (n = 6), FLAG-Y322F DNAM-1 (n = 3), or FLAG-G307S-Y322F DNAM-1 (n = 3) after stimulation with anti-CD3 mAb and either anti-FLAG mAb or control Ig. Data are representative of two independent experiments (**A**) and independent experiments by using Tconv cells derived from three (**B and C**) and two (**D**) different healthy donors. Statistical analysis was performed by using two-way ANOVA with Tukey multiple comparisons test (**B-D**); *P<0.05, **P<0.01, ***P<0.001, ns, not significant.

Next, we established human Tconv-cell transfectants stably expressing FLAG-Y322F DNAM-1 or FLAG-G307S-Y322F DNAM-1, in addition to those expressing FLAG-WT DNAM-1 or FLAG-G307S DNAM-1 (Figure 2-figure supplement 1B). After stimulation with anti-CD3 and anti-FLAG mAbs, Tconv cells expressing FLAG-G307S DNAM-1 produced significantly larger amounts of IFN-γ and TNF and showed significantly greater BrdU incorporation than did those expressing FLAG-WT DNAM-1 (Figure 2B-D). In contrast, production of IFN-γ and TNF and BrdU incorporation were comparable between Tconv cells expressing FLAG-WT and those expressing FLAG-G307S DNAM when the cells were stimulated with anti-CD3 mAb alone. Moreover, enhanced cytokine production and BrdU incorporation were not observed in FLAG-G307S-Y322F DNAM-1-expressing Tconv cells, even after stimulation with anti-CD3 and anti-FLAG mAbs (Figure 2B-D). Taken together, these results suggested that G307S DNAM-1-mediated costimulatory signaling enhanced the tyrosine phosphorylation, pro-inflammatory cytokine production, and proliferation of CD4^+^ Tconv cells.

### G307S mutation enhances recruitment of Lck

To address how the G307S mutation enhances the tyrosine phosphorylation of DNAM-1, we used BioID (proximity-dependent biotin identification) (Roux et al., 2012) to analyze the association of DNAM-1 with Src-family tyrosine kinases such as Lck and Fyn, which are responsible for the tyrosine phosphorylation of DNAM-1 (Shibuya et al., 1999; Zhang et al., 2015, Braun et al., 2020). We established Jurkat transfectants stably expressing FLAG-WT or FLAG-G307S DNAM-1 conjugated with the R118G-mutant BirA (BirA*) at the C-terminus via Gly-Ser linker (Figure 3A). These transfectants expressed comparable levels of FLAG-tagged DNAM-1 (Figure 3B). Although biotinylated molecules were expressed at similar levels in Jurkat transfectants expressing BirA*-conjugated FLAG-WT and those expressing FLAG-G307S DNAM-1 after culture in the presence of biotin (Figure 3-figure supplement 1), transfected cells expressing BirA*-conjugated FLAG-G307S DNAM-1 had more biotinylated Lck than did those expressing BirA*-conjugated FLAG-WT DNAM-1 (Figure 3C). In contrast, biotinylated Fyn was not detected in either transfectant (Figure 3C). Moreover, we found that the enhanced tyrosine phosphorylation of FLAG-G307S DNAM-1 was diminished by treatment with PP2, a selective inhibitor of Src-family kinases (Figure 3D). These results suggested that the G307S mutation augmented the recruitment of Lck to DNAM-1, likely enhancing the tyrosine phosphorylation of DNAM-1.

**Figure 3.**
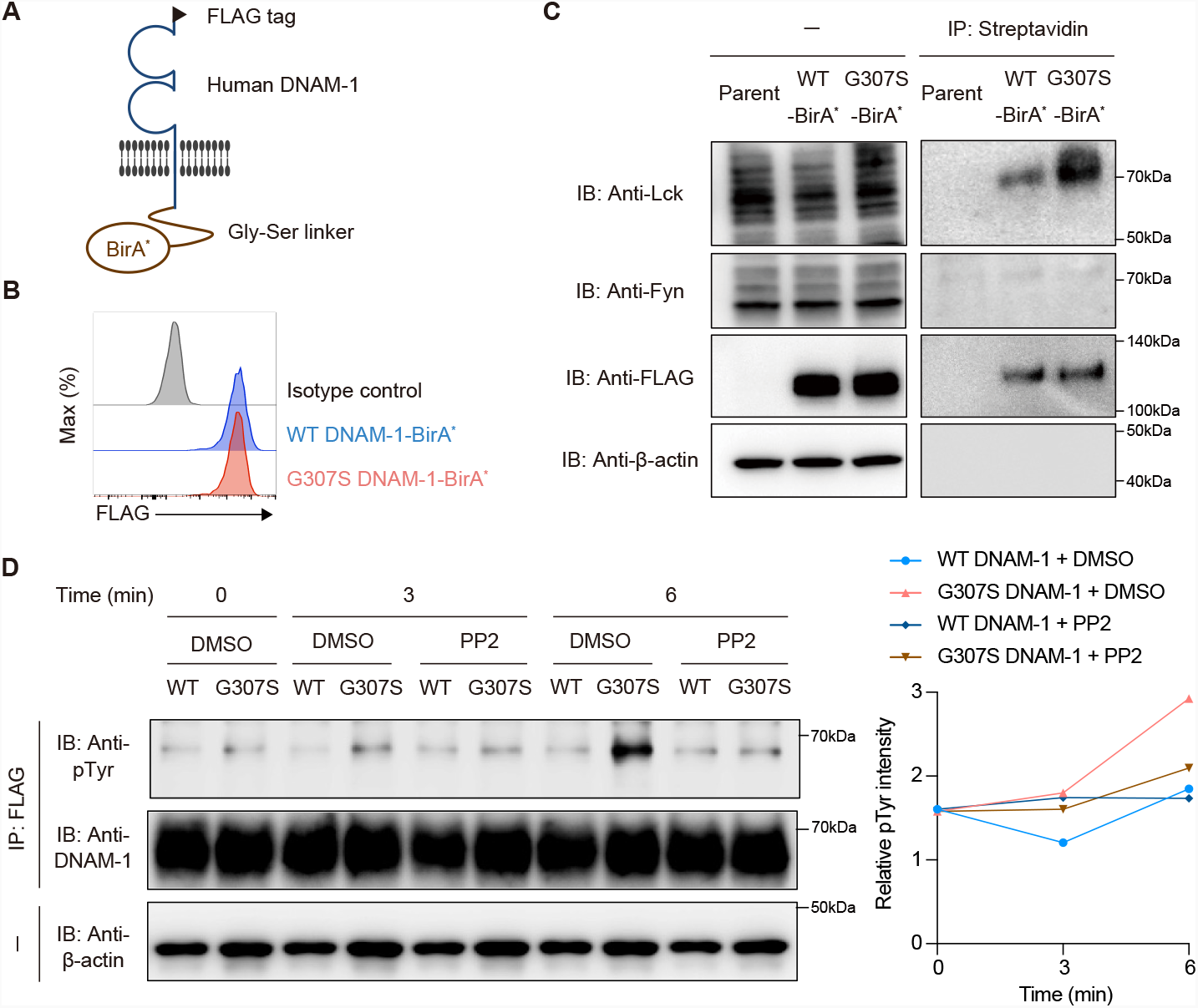
G307S mutation enhanced recruitment of Lck to and tyrosine phosphorylation of DNAM-1. (**A**) Schematic diagram of FLAG-DNAM-1-BirA^*^ fusion protein. (**B**) Expression of FLAG-DNAM-1-BirA^*^ on Jurkat transfectants. (**C**) Jurkat T cells expressing FLAG-WT DNAM-1-BirA^*^ or FLAG-G307S DNAM-1-BirA^*^ were cultured in the presence of biotin, immunoprecipitated with streptavidin, and immunoblotted with ant-Lck, anti-Fyn, anti-FLAG, or anti-β-actin mAbs. (**D**) Jurkat T cells expressing FLAG-WT DNAM-1 or FLAG-G307S DNAM-1 were treated with 25 μM PP2, then stimulated with anti-DNAM-1 mAb, immunoprecipitated or not with anti-FLAG mAb, and immunoblotted with anti-phosphotyrosine, anti-DNAM-1, or anti-β-actin mAbs. Data are representative of three independent experiments (**C and D**).

### G307S mutation of DNAM-1 exacerbates EAE

We next examined the effect of the G307S mutation in CD4^+^ Tconv cells on EAE in mice. Because the glycine at residue 307 of human DNAM-1 is not conserved in mouse DNAM-1 (Tahara-Hanaoka et al., 2005), we generated chimeric DNAM-1 (chDNAM-1) consisting of the extracellular, transmembrane, and a part of intracellular regions (residues 276 to 285) of mouse DNAM-1 fused with the part of the intracellular region (residues 286 to 336) of human WT or G307S DNAM-1 (Figure 4A). We established transfectants of CD4^+^ T cells, which were derived from DNAM-1-deficient myelin oligodendrocyte glycoprotein (MOG) peptide-specific 2D2 TCR transgenic mice (Figure 4B). These transfectants stably expressed WT chDNAM-1 or G307S chDNAM-1 or were mock DNAM-1-deficient 2D2 cells (Figure 4-figure supplement 1). When the mock or WT chDNAM-1-expressing CD4^+^ T cells were co-cultured with CD11c^+^ splenocytes in the presence of 10 µg/mL MOG_35–55_ peptides, the WT chDNAM-1-expressing 2D2 CD4^+^ T cells produced significantly greater amounts of IFN-γ and TNF than did the mock CD4^+^ T cells (Figure 4C), indicating that DNAM-1 plays an important role in the production of Th1 cytokines.

**Figure 4.**
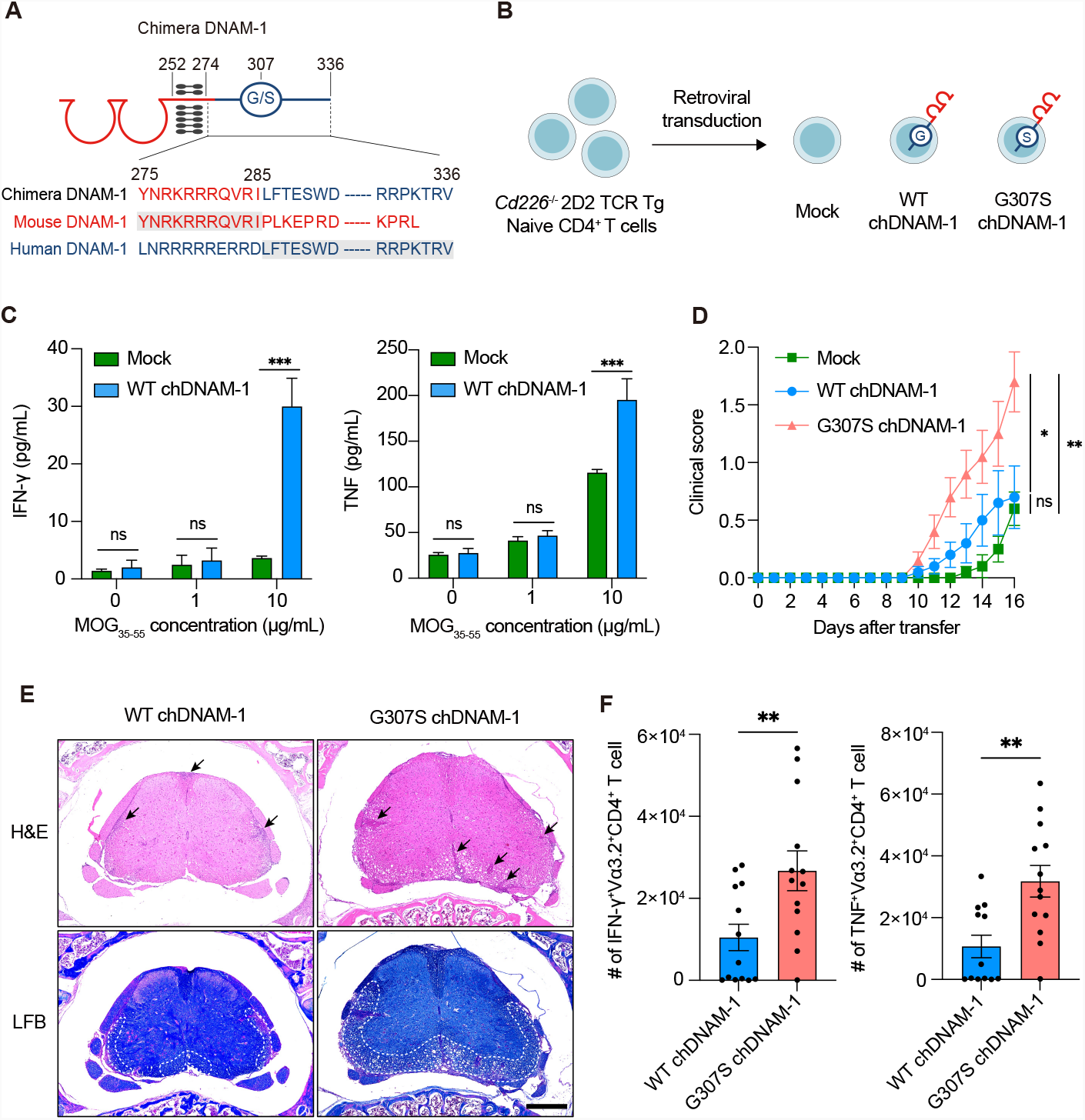
G307S mutation of DNAM-1 exacerbated experimental autoimmune encephalomyelitis. (**A**) Schematic diagram of chimeric DNAM-1 (chDNAM-1) of human WT or G307S DNAM-1. (**B**) Naïve CD4^+^ T cells from *Cd226*^*-/-*^ 2D2 TCR Tg mice were transduced with retroviruses encoding Mock, WT, or G307S chDNAM-1. (**C**) The concentration of IFN-γ and TNF in the culture supernatant of 2D2 CD4^+^ T cells expressing Mock or WT chDNAM-1 after co-culturing with CD11c^+^ splenocytes in the presence of MOG_35-55_ peptides and LPS. (n = 3 in each group) (**D**) Clinical score of EAE in mice that received 2D2 Th1 cells expressing Mock, WT, or G307S chDNAM-1. (n = 10 in each group) (**E**) Representative images of spinal cord sections stained with hematoxylin and eosin (H&E) and luxol fast blue (LFB) from mice that received 2D2 CD4^+^ Th1 cells expressing WT chDNAM-1 or G307S chDNAM-1 at day 21 post-adoptive cell transfer. Black arrows indicate inflammatory cell infiltration, and the demyelination area is shown within white dashed lines. Scale bar: 400 μm. (**F**) Number of IFN-γ-(left) and TNF-(right) expressing TCR Vα3.2^+^ 2D2 CD4^+^ T cell in the spinal cord at day 16 post-adoptive cell transfer. (WT chDNAM-1, n = 13; G307S chDNAM-1, n = 13) Data are representative of three independent experiments (**C**) and pooled of two independent experiments (**D**) and of three independent experiments (**F**) Statistical analysis was performed by using two-way ANOVA with Tukey multiple comparisons test (**D**) and unpaired *t-*test (**C and F**); *P<0.05, **P<0.01, ***P<0.001, ns, not significant.

To clarify the role of G307S DNAM-1-expressing autoreactive CD4^+^ T cells in the pathogenesis of EAE, 2D2 Th1 transfectants expressing Mock, WT chDNAM-1, or G307S chDNAM-1 were polarized into Th1 cells by the culture in the presence of IL-2 and IL-12 and adoptively transferred into irradiated mice in which EAE was then induced by injecting pertussis toxin. Mice that received the 2D2 Th1 transfectant expressing G307S chDNAM-1 had significantly higher clinical scores after cell transfer than did those that received the 2D2 Th1 transfectant expressing WT chDNAM-1 (Figure 4D). Histological analysis by hematoxylin and eosin (H&E) staining and luxol fast blue (LFB) staining demonstrated that inflammatory cell infiltration and demyelination were more severe in mice that received the 2D2 Th1 transfectant expressing G307S chDNAM-1 than in those that received the 2D2 Th1 transfectant expressing WT chDNAM-1 on day 21 after cell transfer (Figure 4E). Furthermore, in the spinal cord, the numbers of IFN-γ^+^ and TNF^+^ transfectants expressing G307S chDNAM-1 were significantly larger than those expressing WT chDNAM-1 on day 16 after cell transfer (Figure 4F, Figure 4- figure supplement 2 A-C). Together, these results indicated that autoreactive 2D2 CD4^+^ T cells expressing G307S chDNAM-1 exacerbated EAE, unlike those expressing WT chDNAM-1.

## Discussion

We showed here that the G307S mutation enhanced proinflammatory cytokine production by CD4^+^ T cells. We further showed that G307S enhanced the recruitment of Lck to, and the tyrosine phosphorylation of, DNAM-1, consistent with a previous report that phosphorylation of the DNAM-1 downstream signaling molecules VAV1 and extracellular signal-regulated kinase (ERK) is increased in rs763361 carriers compared with non-carriers (Gaud et al., 2018). However, it remains unclear how the G307S mutation enhances the recruitment of Lck to DNAM-1. One possible mechanism is that the G307S mutation of DNAM-1 may alter hydrogen bonding, thus affecting the three-dimensional structure of DNAM-1 and protein-protein interaction (Whiteman et al., 1998; Qiu et al., 2018), and leading to the increased binding affinity of Lck for DNAM-1. Future structural analyses of the intracellular region and Lck may be required.

DNAM-1 is expressed on NK cells (Shibuya et al., 1996; Zhang et al., 2015), CD8^+^ T cells (Iguchi-Manaka et al., 2008; Nabekura et al., 2010), innate lymphoid cells (Campana et al., 2019; Nabekura et al., 2020), B cells (Nagayama-Hasegawa et al., 2020), and dendritic cells (Tahara-Hanaoka et al., 2006) in addition to CD4^+^ T cells, which also contribute to the pathogenesis of inflammatory diseases. In particular, DNAM-1 expression on CD8^+^ T cells and B cells, as well as on CD4^+^ T cells, is higher in patients with autoimmune diseases than in healthy controls (Ayano et al., 2015; Nakano et al., 2021; Eschborn et al., 2021). Therefore, G307S DNAM-1-expressing immune cells other than CD4^+^ T cells might also contribute to the development of autoimmune diseases.

rs763361 is associated with various autoimmune disorders, and its minor allele frequency was over 40% in analyzed populations (Bai et al., 2020). Treatment with an anti-DNAM-1 neutralizing mAb has ameliorated several inflammatory disorders in mice (Sato et al., 2021; Dardalhon et al., 2005; Nabekura et al., 2010). Because our results suggest that G307S DNAM-1 contributes more than WT DNAM-1 to the development of inflammatory responses by CD4^+^ T cells, administration of an anti-DNAM-1 neutralizing mAb would more effectively ameliorate inflammatory diseases in rs763361 carriers than non-carriers.

## Materials and Methods

### Mice

WT C57BL/6 mice were purchased from CLEA Japan (Tokyo, Japan). 2D2 TCR Transgenic (Tg) mice (Bettelli et al., 2003) were purchased from Jackson Laboratories (Bar Harbor, ME). DNAM-1-deficient (*Cd226*^*−/−*^) mice on a C57BL/6 background were generated in our laboratory (Iguchi-Manaka et al., 2008). *Cd226*^*−/−*^ mice were crossed with 2D2 TCR Tg mice to generate *Cd226*^*−/−*^ 2D2 TCR Tg mice. These mice were housed under specific-pathogen-free conditions in the same room in the Animal Resource Center at the University of Tsukuba. All animal experiments in this study were performed humanely after receipt of approval from the Animal Ethics Committee of the Laboratory Animal Resource Center, University of Tsukuba and in accordance with the “Fundamental Guideline for Proper Conduct of Animal Experiment and Related Activities in Academic Research Institutions” under the Jurisdiction of the Ministry of Education, Culture, Sports, Science, and Technology.

### Plasmid construction

To generate human DNAM-1 expression vectors, cDNAs encoding WT and G307S DNAM-1 were obtained from human PBMCs, tagged at the 5ʹ-end with cDNA encoding FLAG peptide (DYKDDDDK), and inserted into pMXs-IRES-GFP expression vector (Cell Biolabs, San Diego, CA). To generate Y322F and G307S-Y322F DNAM-1, Phe^322^ was substituted for the Tyr^322^ residue of WT and G307S DNAM-1 by using QuikChange II Site-Directed Mutagenesis Kits (Agilent technologies, Palo Alto, CA) with primers (5′-GAGAGGATATTTTTGTCAACTATCCAACC-3′ and 5′-GGTTGGATAGTTGACAAAAATATCCTCTC-3′). To generate BirA-conjugating human DNAM-1 expression vectors, cDNAs encoding full-length human WT and G307S DNAM-1 tagged with the FLAG peptide at the N-terminus were connected with a Gly-Ser linker (GGGGSGGGGS) followed by cDNA encoding BirA* at the 3ʹ-end by overlap extension PCR with primers (5′-GTGGATCCGCCGCCACCATGTCTGCACTTCTGATCCTAGCTCTTGTTG-3′, 5′-GGAACCTCCGCCCCCGCTCCCTCCGCCACCAACTCTAGTCTTTGGTCTGCGAGAG-3′, 5′-GGTGGCGGAGGGAGCGGGGGCGGAGGTTCCAAGGACAACACCGTGCCCCTGAAGC -3′, and 5′-ACGTCGACTTACTTCTCTGCGCTTCTCAGGGAGATTTC-3′), and inserted into pCDH expression vector (System Biosciences, Mountain View, CA). To generate chDNAM-1-expressing vectors, cDNAs encoding full-length mouse DNAM-1 were restricted by EcoRI, the restriction site of which is in the intracellular region. The N-terminus portion of EcoRI-restricted mouse DNAM-1 was ligated with the PCR products of human WT or G307S DNAM-1 amplified with primers (5′-CTCTGAATTCTATTTACAGAGTCCTGG-3′ and 5′-CTCTGCGGCCGCTTAAACTCTAGTCTTTG-3′). The ligation products were inserted into pMX retroviral expression vector (Cell Biolabs).

### Establishment of transfectants

293GP packaging cells were transfected with retroviral expression vectors by using Lipofectamine 3000 (Thermo Fisher Scientific, Waltham, Massachusetts) in accordance with the manufacturer’s instructions. The medium was replaced 16 h after transfection, and the retroviral supernatant was harvested 3 and 5 days after transfection. After filtration through a 0.45-μm filter (Advantec, Tokyo, Japan), the retrovirus supernatant was concentrated by centrifugation at 5000g at 4 °C for 2 h with Amicon Ultra-15 Centrifugal Filter Units (Millipore, Billerica, MA). A human T cell line (Jurkat) and a mouse T cell line (BW5147) were cultured in complete RPMI-1640 culture medium (Sigma-Aldrich, St. Louis, MO) [10% fetal calf serum (Thermo Fisher Scientific), 50 μM 2-mercaptoethanol, 2 mM L-glutamine, 100 U/mL penicillin, 0.1 mg/mL streptomycin (Sigma-Aldrich), 10 mM HEPES, 1 mM sodium pyruvate, and 100 μM MEM non-essential amino acids (Thermo Fisher Scientific)]. To generate Jurkat transfectants, culture of Jurkat T cells was treated with culture supernatants containing retroviruses encoding FLAG-WT DNAM-1, FLAG-G307S DNAM-1, FLAG-Y322F DNAM-1, FLAG-G307S-Y322F DNAM-1, FLAG-WT DNAM-1-BirA*, or FLAG-G307S DNAM-1-BirA* in the presence of 10 μg/mL protamine sulfate (Sigma-Aldrich) and centrifuged at 1000g at 32 °C for 60 min. FLAG^+^ cells were sorted by flow cytometry, and the equivalent expressions of FLAG and DNAM-1 were confirmed.

### Immunoblotting

To analyze tyrosine phosphorylation of DNAM-1, FLAG-DNAM-1-expressing Jurkat transfectants were incubated with 10 µg/mL anti-human DNAM-1 mAb (clone TX94) (Okumura et al., 2017) on ice for 30 min, followed by 15 µg/mL rabbit polyclonal antibody against mouse IgG (Bethyl Laboratories, Montgomery, TX) for 0, 3, 6, or 10 min at 37 °C. The cells were then lysed with 1% NP-40 lysis buffer (150 mM NaCl, 50 mM Tris, pH 8.0) supplemented with protease inhibitors (1 mM PMSF, 20 U/mL aprotinin) and phosphatase inhibitors (1 mM EGTA, 10 mM NaF, 1 mM Na_4_PO_7_, 0.1 mM β-glycerophosphate, 1 mM Na_3_VO_4_). After incubation for 1 h on ice, the cell lysates were centrifuged at 13,000g for 10 min at 4 °C and immunoprecipitated with anti-FLAG M2 affinity gel (Sigma-Aldrich). Immunoprecipitates were resolved by SDS-PAGE, transferred onto polyvinylidene difluoride (PVDF) membranes by electroblotting, and immunoblotted with HRP-conjugated anti-pTyr Ab (clone 4G10, Millipore) and biotinylated anti-human DNAM-1 mAb (clone TX94) followed by HRP-conjugated streptavidin. In some experiments, FLAG-DNAM-1-expressing Jurkat transfectants were incubated with 25 μM PP2 (Millipore) for 30 min at 37 °C before crosslinking with secondary antibodies.

To analyze the biotinylated proteins induced by FLAG-DNAM-1-BirA*, FLAG-DNAM-1-BirA*-expressing Jurkat transfectants were cultured for 12 h at 37 °C in complete RPMI-1640 supplemented with 0.2-or 1-mM D-biotin (Nakalai Tesque, Kyoto, Japan). Cells were lysed with RIPA lysis buffer [50 mM Tris-HCl (pH 7.5), 150 mM NaCl, 1% NP-40, 0.5% sodium deoxycholate, 2 mM MgCl_2_] supplemented with cOmplete Protease Inhibitor Cocktail (Roche, Basel, Switzerland). After 15 min of incubation on ice, cell lysates were centrifuged at 13,000g for 10 min at 4 °C and immunoprecipitated with Dynabeads MyOne Streptavidin C1 (Invitrogen, Carlsbad, CA). The immunoprecipitated samples were transferred onto PVDF membranes as described earlier and immunoblotted with HRP-conjugated anti-Fyn mAb (clone 15, Santa Cruz Biotechnology, Santa Cruz, CA) and anti-Lck mAb (clone 3A5, Santa Cruz Biotechnology), followed by HRP-conjugated anti-mouse IgG Ab, HRP-conjugated anti-FLAG Ab (clone M2, Sigma-Aldrich), and HRP-conjugated anti-β-actin mAb (clone 8H10D10, Cell Signaling Technology, Beverly, MA).

### Passive EAE model

Naïve CD4^+^ T cells were purified from CD4^+^ T cells from pooled spleens and lymph nodes of *Cd226*^*−/−*^ 2D2 TCR Tg mice by using mouse CD4 MicroBeads (Miltenyi Biotec, Bergisch-Gladbach, Germany), followed by flow cytometric sorting for CD25^-^CD44^Lo^CD62L^Hi^CD4^+^ cells. The purity was greater than 99%. Sorted naïve 2D2 CD4^+^ T cells were activated with plate-coated 2 μg/mL anti-mouse CD3e mAb (clone 145-2C11, BioLegend, San Diego, CA) and soluble 1 μg/mL anti-mouse CD28 mAb (clone 37.51, BioLegend) under Th1-polarizing conditions [1 ng/mL mouse IL-2 (BD Bioscience, San Diego, CA) and 5 ng/ml mouse IL-12 (BD Biosciences)] in complete RPMI-1640 culture medium. On day 2, retrovirus supernatant was added in the presence of 10 μg/mL protamine sulfate (Sigma-Aldrich) and the cell mixture was centrifuged at 1000g at 32 °C for 60 min. Five days after initial stimulation, the retrovirally transduced 2D2 CD4^+^ T cells were intravenously injected into 4-Gy-irradiated WT C57BL/6 mice (5 × 10^5^ cells/mouse). Pertussis toxin 200 ng (#180, List Labs, Campbell, CA) in 100 μL PBS was injected intraperitoneally before, and 2 days after, the T cell transfer. In all experiments, the equivalent expression of T-bet, IFN-γ, and DNAM-1 was confirmed by flow cytometry. Mice were monitored daily for the development of EAE in accordance with the criteria provided by Hooke Laboratories (https://hookelabs.com/protocols/eaeAI_C57BL6.html). The number of the total cells isolated from the spinal cord was counted by TC20 Automated Cell Counter (Bio-Rad, Hercules, CA), and the number of spinal cord-infiltrating TCR Vα3.2^+^ 2D2 CD4^+^ T cells and TCR Vα3.2^-^ non-2D2 CD4^+^ T cells were calculated based on the percentage obtained by flow cytometry.

### Histological analysis

The spinal cord on day 21 after adoptive cell transfer was fixed with 4% paraformaldehyde, embedded in paraffin, and sectioned. Fixed sections were stained with hematoxylin and eosin (H&E) and luxol fast blue (LFB) according to standard protocols and observed via microscopy.

### Co-culture assay of 2D2 CD4^+^ T cells and CD11c^+^ cells

Splenic CD4^+^ T cells were purified from *Cd226*^*−/−*^ 2D2 TCR transgenic mice by using mouse CD4 MicroBeads (Miltenyi Biotec). Sorted 2D2 CD4^+^ T cells were activated with plate-coated 2 μg/mL anti-mouse CD3e mAb (clone 145-2C11, BioLegend), soluble 1 μg/mL anti-mouse CD28 mAb (clone 37.51, BioLegend), and 1 ng/mL mouse IL-2 (BD Biosciences) in complete RPMI-1640 culture medium. On day 2, retrovirus supernatant was added in the presence of 10 μg/mL protamine sulfate (Sigma-Aldrich) and the cells were centrifuged at 1000g at 32 °C for 60 min. At 6 days after initial stimulation, CD11c^+^ cells were sorted from pooled spleens of WT C57BL/6 mice by using mouse CD11c MicroBeads (Miltenyi Biotec). Then, 1 × 10^4^ sorted CD11c^+^ splenocytes were co-cultured for 48 h at 37 °C with 5 × 10^4^ retrovirally transduced 2D2 CD4^+^ T cells in the presence of 10 μg/mL MOG_35–55_ peptides (MBL, Boston, MA) and 25 ng/mL LPS.

### Retroviral transduction of human Tconv cells and Treg cells

Peripheral blood was obtained from eight healthy Japanese volunteers (average age, 30.4 years; minimum, 26 years; maximum, 40 years) after written informed consent had been obtained, along with approval by the Tsukuba Clinical Research & Development Organization (T-CReDO) of the University of Tsukuba. All blood samples were collected by heparin blood sampling. Human samples were de-identified before use. Human PBMCs were isolated by using Lymphoprep (Stemcell Technologies, Vancouver, BC, Canada) gradient centrifugation. CD4^+^ T cells were purified from human PBMCs by using human CD4 MicroBeads (Miltenyi Biotec). CD4^+^ CD25^Lo^ CD127^Hi^ Tconv cells and CD4^+^ CD25^Hi^ CD127^Lo^ Treg cells were sorted by flow cytometry and stimulated in 96-well cell-culture plates with three times the number of Dynabeads M-450 Tosylactivated beads (Thermo Fisher Scientific) coated with anti-human CD3 mAb (clone HIT3α, BD Biosciences) (50%) and anti-human CD28 mAb (clone CD28.2, BD Biosciences) (50%) in complete RPMI-1640 culture medium supplemented with 10 ng/mL human IL-2 (BD Biosciences). On day 1, retrovirus supernatant was added into the culture medium of Tconv cells or Treg cells in the presence of 10 μg/mL protamine sulfate and the mixture was centrifuged at 850g at 32 °C for 90 min. On day 5, the anti-human CD3 mAb-and anti-human CD28 mAb-coated beads were removed, and the culture medium was replaced with a complete RPMI-1640 culture medium supplemented with 10 ng/mL human IL-2. On day 7, GFP^+^ retrovirally transduced cells were sorted by flow cytometry and stimulated in 96-well cell-culture plates with three times the number of anti-human CD3 mAb-and anti-human CD28 mAb-coated beads in complete RPMI-1640 culture medium supplemented with 10 ng/mL human IL-2. On day 12, the anti-human CD3 mAb-and anti-human CD28 mAb-coated beads were removed, and the culture medium was replaced with complete RPMI-1640 culture medium supplemented with 0.1 ng/mL human IL-2. On day 14, FLAG, DNAM-1, and Foxp3 expression was analyzed by flow cytometry.

### Functional assay of Tconv cells and Treg cells

To analyze cytokine production by Tconv cells, FLAG-DNAM-1-expressing human Tconv cells were stimulated for 12 h with plate-bound 0.25 μg/mL anti-human CD3 mAb (clone HIT3α, BD Biosciences) and either plate-bound 3 μg/mL mouse IgG1 isotype control or anti-FLAG mAb (clone M2, Sigma-Aldrich) in complete RPMI-1640 culture medium. The culture supernatants were harvested, and cytokine concentrations were analyzed by BD Cytometric Bead Array (BD Biosciences). To analyze cell proliferation, FLAG-DNAM-1-expressing human Tconv cells were stimulated for 24 h with plate-bound 0.25 μg/mL anti-human CD3 mAb (clone HIT3α, BD Biosciences) and either plate-bound 3 μg/mL mouse IgG1 isotype control or anti-FLAG mAb (clone M2, Sigma-Aldrich) in complete RPMI-1640 culture medium supplemented with 10 μM BrdU. In accordance with the manufacturer’s instructions, the cells were stained for BrdU by using an APC BrdU Flow Kit (BD Biosciences) and analyzed by flow cytometry. To analyze Foxp3 expression and IL-10 production by Treg cells, FLAG-DNAM-1-expressing human Treg cells were stimulated in complete RPMI-1640 culture medium at 37 °C for 5 days with plate-bound 0.2 or 1 μg/mL anti-human CD3 mAb (clone HIT3α) and either plate-bound 30 μg/mL mouse IgG1 isotype control or anti-FLAG mAb (clone M2, Sigma-Aldrich). The culture supernatants were harvested, and cytokine concentrations were analyzed by BD Cytometric Bead Array.

### *In vitro* Treg suppression assay

FLAG-DNAM-1-expressing human Treg cells were co-cultured in complete RPMI-1640 culture medium with 2 × 10^4^ CellTrace Violet (Thermo Fisher Scientific)-labeled Tconv cells (freshly isolated from the same donor as that of the Treg cells) at 0.5: 1, 0.25: 1, or 0.125: 1 in the presence of 4 × 10^4^ Dynabeads M-450 Tosylactivated beads coated with anti-human CD3 mAbs (clone HIT3α) (10%) and anti-FLAG mAbs (clone M2) (90%). The cells were incubated at 37 °C for 96 h, and then the CellTrace Violet signals were analyzed by flow cytometry. Suppression (%) was calculated by using the formula: [100 – (% division of Tconv + Treg) / (% division of stimulated Tconv alone) × 100].

### Measurement of cytokines in culture supernatant

Concentrations of IFN-γ, TNF, IL-4, IL-17A, and IL-10 in the culture supernatants of stimulated mouse CD4^+^ T cells, human Tconv cells, or Treg cells were measured with BD Cytometric Bead Array Human Flex Sets for IFN-γ, TNF, IL-4, IL-17A, and IL-10 (BD Biosciences).

### Flow cytometry

Suspensions of human Tconv cells or Treg cells were stained with V500-conjugated anti-human CD4 (clone RPA-T4, BD Biosciences), PE-Cy7-conjugated anti-human CD25 (clone BC96, Biolegend), APC-conjugated anti-human CD127 (clone eBioRDR5, Invitrogen), PE-conjugated anti-DYKDDDDK (clone L5, Biolegend), biotinylated anti-human DNAM-1 (clone TX94), and APC-conjugated streptavidin. The suspension of mouse CD4^+^ T cells and the spinal cord sections were stained with APC-Cy7-conjugated anti-mouse CD4 (clone GK1.5, Biolegend), BV510-conjugated anti-mouse CD4 (clone RM4-5, Biolegend), BB515-conjugated anti-mouse CD25 (clone PC61, BD Biosciences), APC-conjugated anti-mouse CD44 (clone IM7, BD Biosciences), PE-conjugated anti-mouse CD62L (clone MEL-14, BD Biosciences), PE-conjugated anti-mouse TCR Vα3.2 (clone RR3–16, Biolegend), Alexa Fluor 700-conjugated anti-mouse CD45.2 (clone 104, Biolegend), PE-Cy7-conjugated anti-mouse TCR β chain (clone H57-597, Biolegend), BV711-conjugated anti-mouse CD8 (clone 53-6.7, Biolegend), biotinylated anti-mouse/human B220 (clone RA3-6B2, Biolegend), biotinylated anti-mouse NK1.1 (clone PK136, eBioscience, SanDiego, California), biotinylated anti-mouse/human CD11b (clone M1/70, Biolegend), biotinylated anti-mouse CD11c (clone HL3, BD Biosciences), biotinylated anti-mouse Ly-6G/Ly-6C (clone RB6-8C5, Biolegend), biotinylated anti-mouse DNAM-1 (clone TX42.1), and BV605-or APC-conjugated streptavidin. For staining of intracellular IFN-γ and TNF, CD4^+^ T cells were fixed and permeabilized with BD Cytofix/Cytoperm solution (BD Biosciences), washed with Intracellular Staining Perm Wash Buffer (BioLegend), and stained with FITC-conjugated anti-mouse IFN-γ (clone XMG1.2, BD Biosciences) and APC-conjugated anti-mouse TNF (clone MP6-XT22, BD Biosciences). For staining of intracellular human Foxp3 and mouse T-bet cells were fixed and permeabilized by using an eBioscience Foxp3/Transcription Factor Staining Buffer Set (Thermo Fisher Scientific) and stained with Alexa Fluor 647-conjugated anti-human Foxp3 mAb (clone 259D, Biolegend) or PE-Cy7-conjugated anti-mouse T-bet mAb (clone 4B10, Biolegend). In all experiments, double T cells were excluded by FSC-A (forward scatter area) and FSC-H (forward scatter height) gating, followed by SSC-A (side scatter area) and SSC-W (side scatter width) gating. Propidium iodide (PI; Sigma-Aldrich, St. Louis, Missouri, USA) was used to exclude dead cells; living cells were defined as PI-negative cells. In some experiments, cells were stained with Zombie Violet and NIR Fixable Viability Kit (Biolegend); living cells were defined as Zombie Violet-or NIR-negative cells. Samples underwent flow cytometry (LSRFortessa or FACSAria III, Becton Dickinson, Franklin Lakes, New Jersey, USA), and the data were analyzed by using FlowJo software (FlowJo, Ashland, Oregon, USA).

### Statistical analysis

Unpaired two-tailed t-tests and two-way ANOVA were used to compare the data by using Prism 9 (GraphPad Software, La Jolla, California, USA). P < 0.05 was considered statistically significant. Error bars show SEM.

## Author contributions

R. Murata, A. Shibuya, and K. Shibuya designed the research; R. Murata, S. Kinoshita, and K. Matsuda performed the experiments; A. Kawaguchi provided materials for BioID; and R. Murata, A. Shibuya, and K. Shibuya wrote the manuscript. All authors approved the final manuscript and agreed to its publication.

## Acknowledgments

We thank F. Abe, R. Hirochika, and T. Kuroki for technical support; K. Sato, K. Kanemaru, Y. Yamashita-Kanemaru, and T. Nabekura for helpful discussions; and S. Tochihara, W. Saito, M. Kaneko, and H. Furugen for secretarial assistance.

This research was supported by grants provided by the Ministry of Education, Culture, Sports, Science, and Technology of Japan (grant numbers 18H05022 and 21H04836 to A. Shibuya, 21H02708 to K. Shibuya, and 22J11500 to R. Murata), and by JST SPRING (grant number JPMJSP2124 to R. Murata). The sponsors had no control over the interpretation, writing, or publication of this work.

---

**Figure 1-figure supplement 1.**
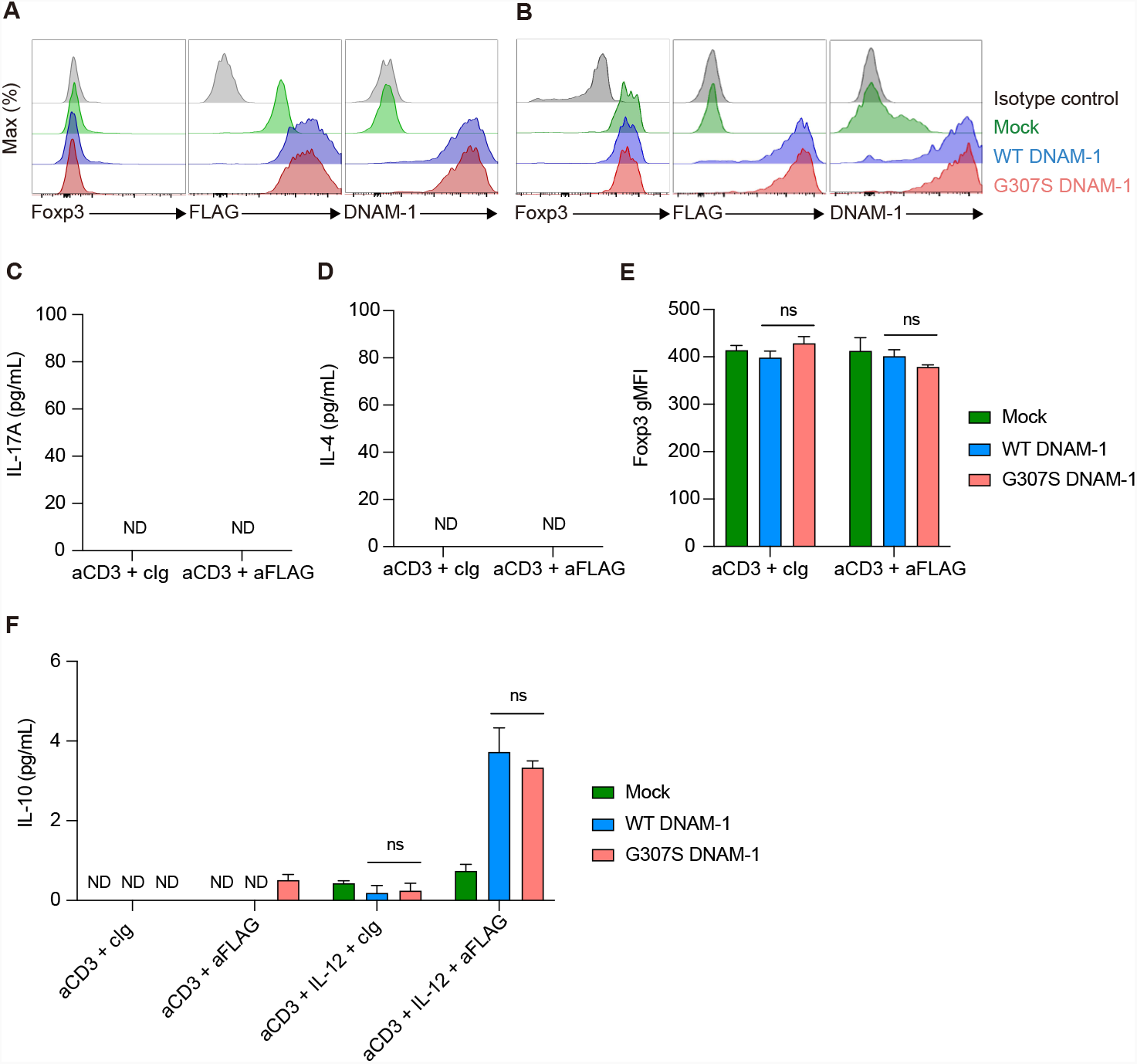
Establishment and functional analysis of FLAG-WT and FLAG-G307S DNAM-1-expressing human CD4^+^ T cells. (**A and B**) Expression of Foxp3, FLAG, and DNAM-1 on human Tconv cells (**A**) and human Treg cells (**B**). (**C and D**) The concentration of IL-17A (**C**) and IL-4 (**D**) in the culture supernatant of human Tconv cells expressing Mock, FLAG-WT DNAM-1, or FLAG-G307S DNAM-1 after stimulation with anti-CD3 mAb and either anti-FLAG mAb or control Ig. (n = 3 in each group) (**E**) The Foxp3 expression in human Treg cells expressing Mock, FLAG-WT DNAM-1, or FLAG-G307S DNAM-1 after stimulation with anti-CD3 mAb and either anti-FLAG mAb or control Ig. (n = 3 in each group) (**F**) The concentration of IL-10 in the culture supernatant of human Treg cells expressing Mock, FLAG-WT DNAM-1, or FLAG-G307S DNAM-1 after stimulation with anti-CD3 mAb and either anti-FLAG mAb or control Ig for five days. (n = 3 in each group) Data are representative of independent experiments using Tconv (**C and D**) and Treg (**E**) cells derived from three healthy donors, and of two independent experiments (**F**). Statistical analysis was performed by using two-way ANOVA with Tukey multiple comparisons test (**C-F**); *P<0.05, **P<0.01, ***P<0.001, ns, not significant, ND. Not detected.

**Figure 2-figure supplement 1.**
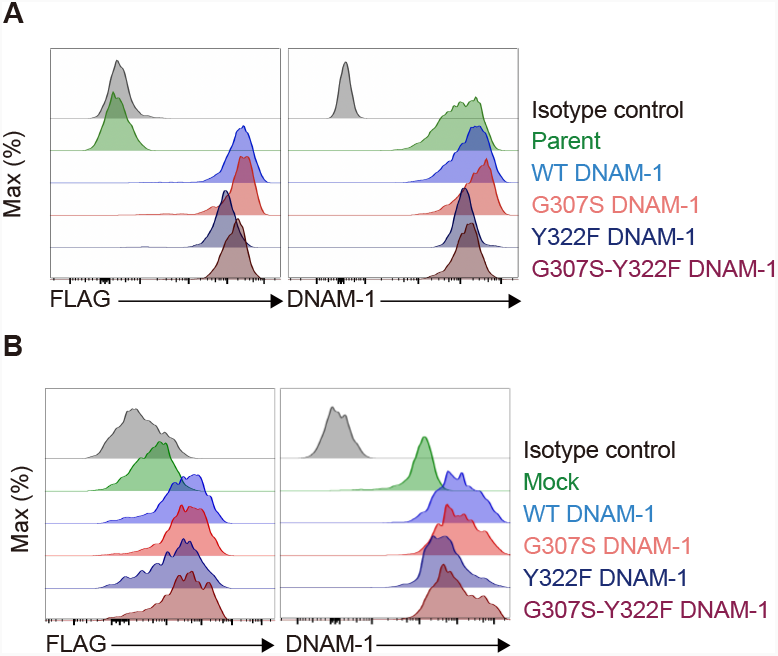
Establishment of FLAG-Y322F and FLAG-G307S-Y322F DNAM-1-expressing Jurkat and human Tconv cells. (**A and B**) Expression of FLAG and DNAM-1 on Jurkat (**A**) and human Tconv (**B**) transfectants. Data are representative of two independent experiments.

**Figure 3-figure supplement 1.**
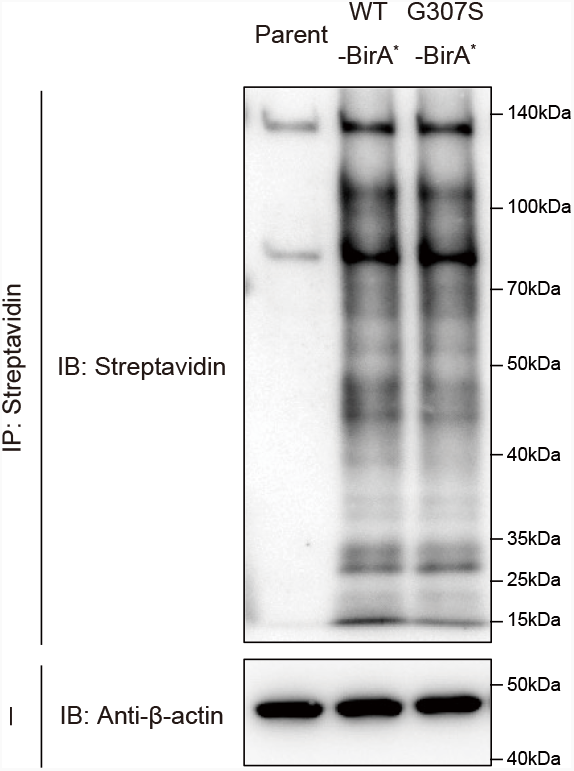
G307S mutation of DNAM-1 doesn’t alter the biotinylation of cytosolic proteins. Jurkat T cells expressing FLAG-WT DNAM-1-BirA^*^ or FLAG-G307S DNAM-1-BirA^*^ were cultured in the presence of biotin, immunoprecipitated or not with streptavidin, and immunoblotted with streptavidin or anti-b-actin mAb. Data are representative of two independent experiments.

**Figure 4-figure supplement 1.**
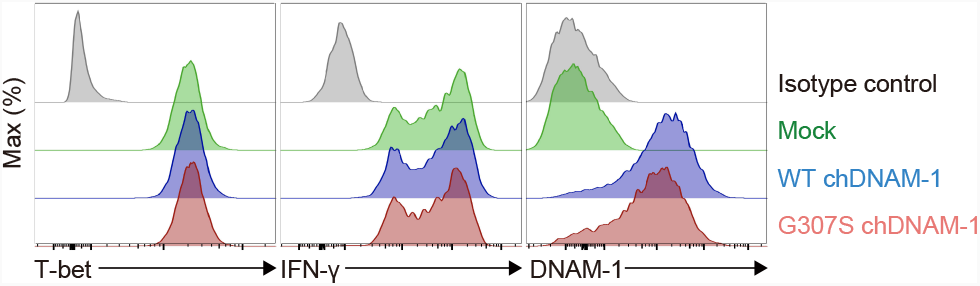
Establishment of chDNAM-1-expressing 2D2 Th1 cells. Expression of T-bet, IFN-γ, and chDNAM-1 in 2D2 CD4^+^ T cells. Data are representative of two independent experiments.

**Figure 4-figure supplement 2.**
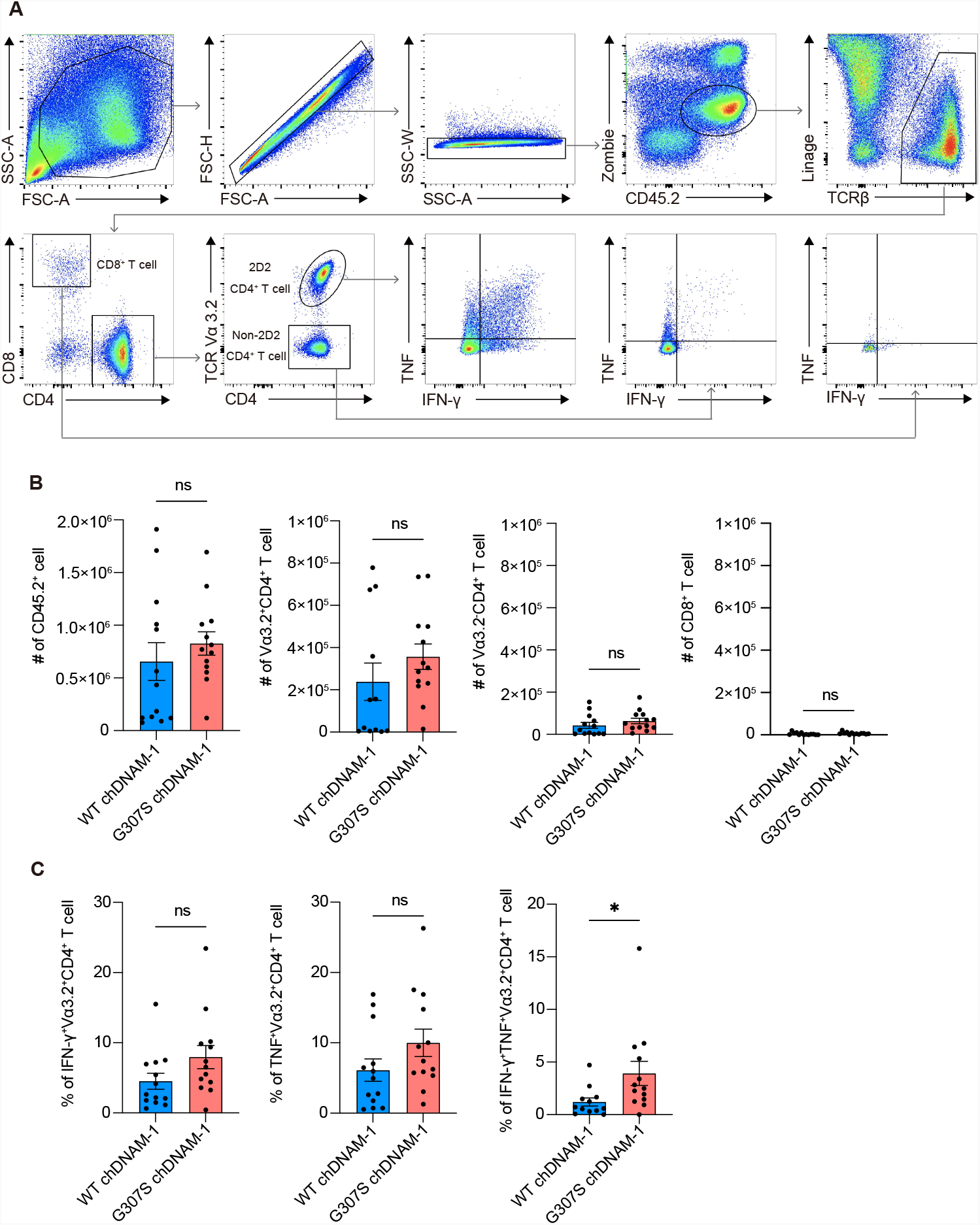
*Ex vivo* analysis of spinal-cord-infiltrating 2D2 CD4^+^ T cells. (**A**) Gating strategy for TCR Vα3.2^+^ 2D2 CD4^+^ T cells, TCR Vα3.2^−^ non-2D2 CD4^+^ T cells, and CD8^+^ T cells in the spinal cord at day 16 post-adoptive cell transfer. Linage includes B220, NK1.1, CD11b, CD11c, and Gr-1. (**B**) Number of CD45.2^+^ cell, TCR Vα3.2^+^ 2D2 CD4^+^ T cell, TCR Vα3.2^−^ non-2D2 CD4^+^ T cell, and CD8^+^ T cell in the spinal cord at day 16 post-adoptive cell transfer. (WT chDNAM-1, n = 13; G307S chDNAM-1, n = 13) (**C**) Percentage of IFN-γ^+^ (left), TNF^+^ (middle), and IFN-γ^+^TNF^+^ (right) TCR Vα3.2^+^ 2D2 CD4^+^ T cell in the spinal cord at day 16 post-adoptive cell transfer. (WT chDNAM-1, n = 13; G307S chDNAM-1, n = 13) Data are representative of three independent experiments (**A**) and pooled of three independent experiments (**B and C**). Statistical analysis was performed by using unpaired *t-*test (**B and C**); *P<0.05, **P<0.01, ***P<0.001, ns, not significant.

